# Absolute measurements of mRNA translation in *C. crescentus* reveal important fitness costs of vitamin B_12_ scavenging

**DOI:** 10.1101/552000

**Authors:** James R. Aretakis, Alisa Gega, Jared M. Schrader

**Author notes:** Corresponding Author Jared M. Schrader Department of Biological Sciences 5047 Gullen Mall Biological Sciences Building Room 2119 Detroit, MI 48202.

## Abstract

*Caulobacter crescentus* is a model for the bacterial cell cycle which culminates in asymmetric cell division, yet little is known about the absolute levels of protein synthesis of the cellular parts needed to complete the cell cycle. Here we utilize ribosome profiling to provide absolute measurements of mRNA translation of the *C. crescentus* genome, providing an important resource for the complete elucidation of the cell cycle gene-regulatory program. Analysis of protein synthesis rates revealed ∼4.5% of cellular protein synthesis are for genes related to vitamin B_12_ import (*btuB*) and B_12_ independent methionine biosynthesis (*metE*) when grown in common growth media lacking B_12_. While its facultative B_12_ lifestyle provides a fitness advantage in the absence of B_12_, we find that it provides lower fitness of the cells in the presence of B_12_, potentially explaining why many *Caulobacter* species have lost the *metE* gene and become obligates for B_12_.

## Introduction

In bacterial systems biology, global mRNA translation measurements are critical for understanding how cells utilize their resources to achieve their evolutionarily selected cell growth and division cycles. To complete the bacterial cell cycle, the protein parts encoded within the genome must be transcribed into mRNAs that are translated into the appropriate number of proteins for the daughter cells to be generated. Genome-wide absolute quantitation of protein level measurements have allowed the monitoring of protein resource allocation (Hui et al., 2015; Li, Burkhardt, Gross, & Weissman, 2014) revealing that these cells allocate resources for optimal growth. As the ribosome content is positively correlated with the growth rate (Bremer & Dennis, 2008), cells must optimize the fraction of protein synthesis needed to make new ribosomes (enzymes that make proteins) vs the protein synthesis needed to produce the proteomes of the daughter cells to achieve fast generation times (Li et al., 2014; Scott, Klumpp, Mateescu, & Hwa, 2014). Optimality has also been observed at the protein-complex level, as translation of a stoichiometric amount of protein subunits to the overall multi-protein complexes has been observed (Li et al., 2014), with different post-transcriptional strategies across species utilized to achieve the optimal protein concentration (Lalanne et al., 2018). Therefore, to understand the mechanisms controlling the growth and division cycles of diverse bacteria, we must understand how bacteria are able to optimize their protein synthesis resources for maximal fitness.

*Caulobacter crescentus* is an oligotrophic α-proteobacteria with a carefully orchestrated cell cycle yielding asymmetric cell division and a model organism for the study of the bacterial cell cycle (Collier, 2016; K. Lasker, Mann, & Shapiro, 2016). In *C. crescentus*, cells undergo changes in gene expression of ∼20% of their entire genome during the process of the cell cycle (Laub, McAdams, Feldblyum, Fraser, & Shapiro, 2000; Schrader et al., 2016). Timing of 57% of the cell cycle-regulated mRNAs are controlled at the transcription level by a master regulatory circuit that is composed of 4 transcription factors and a DNA methylase (K. Lasker et al., 2016; Zhou et al., 2015) and 49% of those cell cycle-regulated mRNAs are additionally regulated at the level of mRNA translation (Schrader et al., 2016). Importantly, global *C. crescentus* studies have focused solely on the control of the timing of gene expression in the cell cycle and thus little is known about the absolute levels of protein synthesis, or how the protein synthesis resources are allocated across the proteome.

Here, we utilize ribosome profiling to achieve a quantitative genome-wide absolute measure of protein synthesis in *C. crescentus*. This resource provides the absolute protein synthesis rate of each protein expressed from the *C. crescentus* genome and a global map of protein synthesis resource allocation. Absolute levels of mRNA translation of cell cycle master-regulators showed higher levels of mRNA translation compared to their known DNA binding sites for all but CcrM, and a relatively low level of mRNA translation of CtrA regulatory proteins relative to the CtrA master regulator itself. PopZ, a polar protein scaffold that recruits asymmetric cell fate specification proteins (Berge & Viollier, 2018), is at a limiting concentration compared to its client proteins suggesting that these clients compete for access to the cell pole. Surprisingly, we discovered that the *btuB* vitamin B_12_ importer and the *metE* methionine biosynthetic gene were among the most highly translated genes in the absence of B_12_, showing that *C. crescentus’* B_12_ scavenging pathway requires a surprisingly large amount of the cells protein synthesis resources. The high cost of protein synthesis of the B_12_ scavenging pathway is reduced in the presence of B_12_ by riboswitches in the 5’ UTR of these two genes. The widely utilized lab strain, NA1000, is a facultative B_12_ scavenger due to the *metE* gene which produces methionine in the absence of B_12_, yet many natural *Caulobacter* isolates are obligate B_12_ scavengers (Poindexter, 1964). We show that while the facultative B_12_ scavenging lifestyle generates a fitness tradeoff, where in the absence of B_12_ there is a positive fitness advantage from MetE’s B_12_-independent methionine production, while in B_12_’s presence there is a fitness disadvantage due to the wasted cost of MetE’s protein synthesis, providing an explanation for why many isolates have lost the *metE* gene to become obligates for B_12_.

## Results

### Absolute quantitation of mRNA translation rates

Ribosome profiling provides a global direct measure of the protein synthesis rate by sequencing ribosome protected mRNA footprints (Ingolia, Ghaemmaghami, Newman, & Weissman, 2009; Li et al., 2014; Oh et al., 2011). To determine absolute rates of translation in *C. crescentus*, ribosome profiling was performed in unsynchronized *C. crescentus* NA1000 cells grown in M2G minimal medium. In fast growing bacteria where the rate of translation is the main driving force of protein levels, protein degradation can be negligible and therefore the main driving force of protein levels is mRNA translation (Li et al., 2014). This is largely true in *C. crescentus*, as >95% of proteins were found to have half-lives longer than the cell cycle (Grunenfelder et al., 2001). First, we examined the ribosome footprint density along each ORF on mRNAs as a relative measure for translation (Fig 1A). For example, in the *divK/pleD* polycistronic operon, we find that *divK* has 2.0 fold higher ribosome density as compared to *pleD* (Fig 1A). For absolute quantitation, it is assumed that the average elongation rate is constant for each mRNA which would allow the average ribosome density to be directly proportional to the rate of protein synthesis of each ORF (Li et al., 2014). It is also assumed that all ribosomes will finish translation and make the full-length protein (Li et al., 2014). Next, to reduce the impact of fast and slow-moving ribosomes in the ribosome occupancy profiles along ORFs on the quantitative level of translation, we used winsorization to correct the average ribosome footprint density of each ORF (Table S1). Start codon and stop codon regions were omitted from the analysis to avoid biases in slow moving ribosomes that are initiating or terminating (Aretakis, Al-Husini, & Schrader, 2018; Oh et al., 2011).

**Figure 1.**
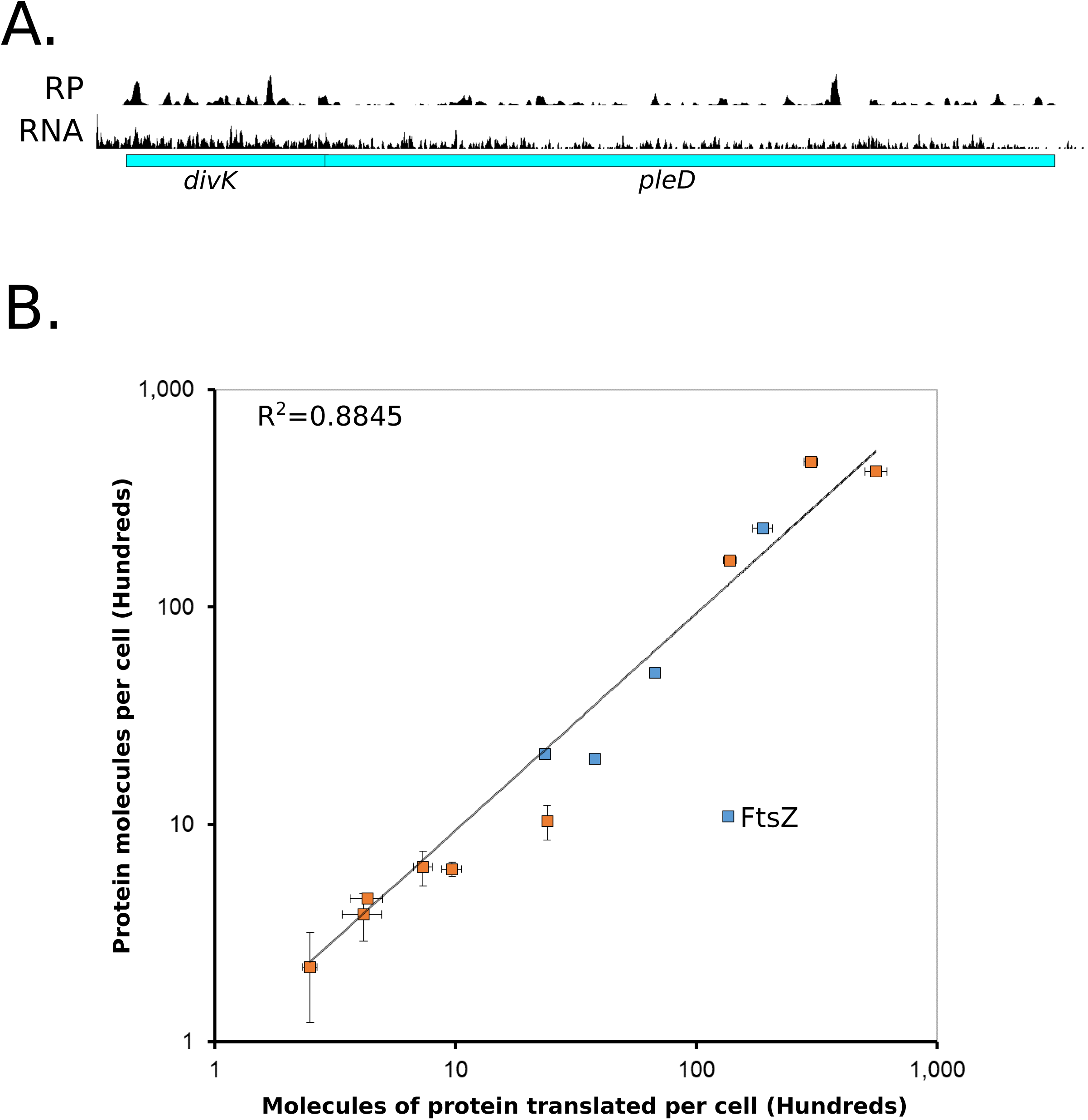
Absolute quantitation of *C. crescentus* protein synthesis by ribosome profiling. A.) Ribosome profiling data for cells grown in M2G media of the *divK/pleD* operon. Average ribosome density of *divK* is 2.0 times higher than for *pleD*. mRNA data from (Schrader et al., 2014). B.) Absolute protein levels of unsynchronized cells measured by western blot (blue) or YFP fusions (orange) compared to the absolute molecules of protein translated per cell calculated by ribosome profiling. Vertical error bars indicate the standard deviation in YFP intensity or standard deviation for the western blots, while horizontal error bars indicate the standard deviation from ribosome profiling replicates (n=3). FtsZ is indicated as its protein levels are under proteolytic control (Kelly et al., 1998). Data in Tables S1 and S2.

To convert ribosome density to absolute mRNA translation rates we measured the average protein mass of *C. crescentus* cells which was multiplied by the fractional ribosome density measurement of each gene and divided by the molecular weight 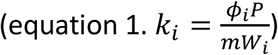 (Li et al., 2014). This measure of the average number of proteins translated per cell correlated well between the protein concentrations reported in the literature as well as protein concentration measurements reported here using YFP intensity of C-terminally tagged gene-fusions (Fig 1B, Fig S1, Table S2). As expected, FtsZ, the key cell division protein which is known to be a substrate of cell cycle dependent proteolysis, (Kelly, Sackett, Din, Quardokus, & Brun, 1998) has a 13-fold higher amount of translated protein than protein observed in the cell, while stable proteins ranged between 0.65 to 2.3-fold. The same phenomenon of higher translation levels than protein levels was also observed for the proteolyzed cell cycle regulators DnaA and CcrM, and upon deletion of the Lon protease which is known to be responsible for their proteolysis the correlation was restored (Peter Chien personal communication, Fig S1) (Jonas, Liu, Chien, & Laub, 2013; Wright, Stephens, Zweiger, Shapiro, & Alley, 1996). These data suggest that the absolute measures of mRNA translation are reflecting the absolute protein synthesis rate for each ORF in *C. crescentus* and provide a reasonable measure of steady state protein levels for stable proteins (data can be found in Table S1).

### Global analysis of *C. crescentus* absolute mRNA translation levels

*C. crescentus* cells dedicate a significant percentage of their protein synthesis to several major cellular processes associated with cell growth (Fig 2A, Fig S2). Across major “KEGG categories” we analyzed the percentage of ribosome footprints to understand their allocation of protein synthesis capacity (Table S3). Nutrient transporters (11.8%), the ribosome (11.2%), and the cell envelope (9.8%) represent the largest classes of protein production for *C. crescentus*. A significant fraction of protein synthesis capacity (25.5%) is allocated to produce proteins of unknown function, showing a significant fraction of the cell’s protein synthesis capacity is not understood. By comparing the fraction of the translation KEGG category across minimal media (M2G) (15.9%) and a richer complex media (PYE) (22.8%) we find the cells dedicate a larger amount of protein synthesis capacity to making translational machinery in rich media similar to *E. coli* (Bremer & Dennis, 2008). Additionally, we see that a larger amount of protein synthesis capacity in “cell growth and death” is observed in M2G (11.3%) compared to PYE (6.45%), owing largely to increased protein synthesis capacity of an operon of cell-contact dependent toxins and immunity proteins that are known to be expressed in stationary phase (Fig S2) (Garcia-Bayona, Guo, & Laub, 2017).

**Figure 2.**
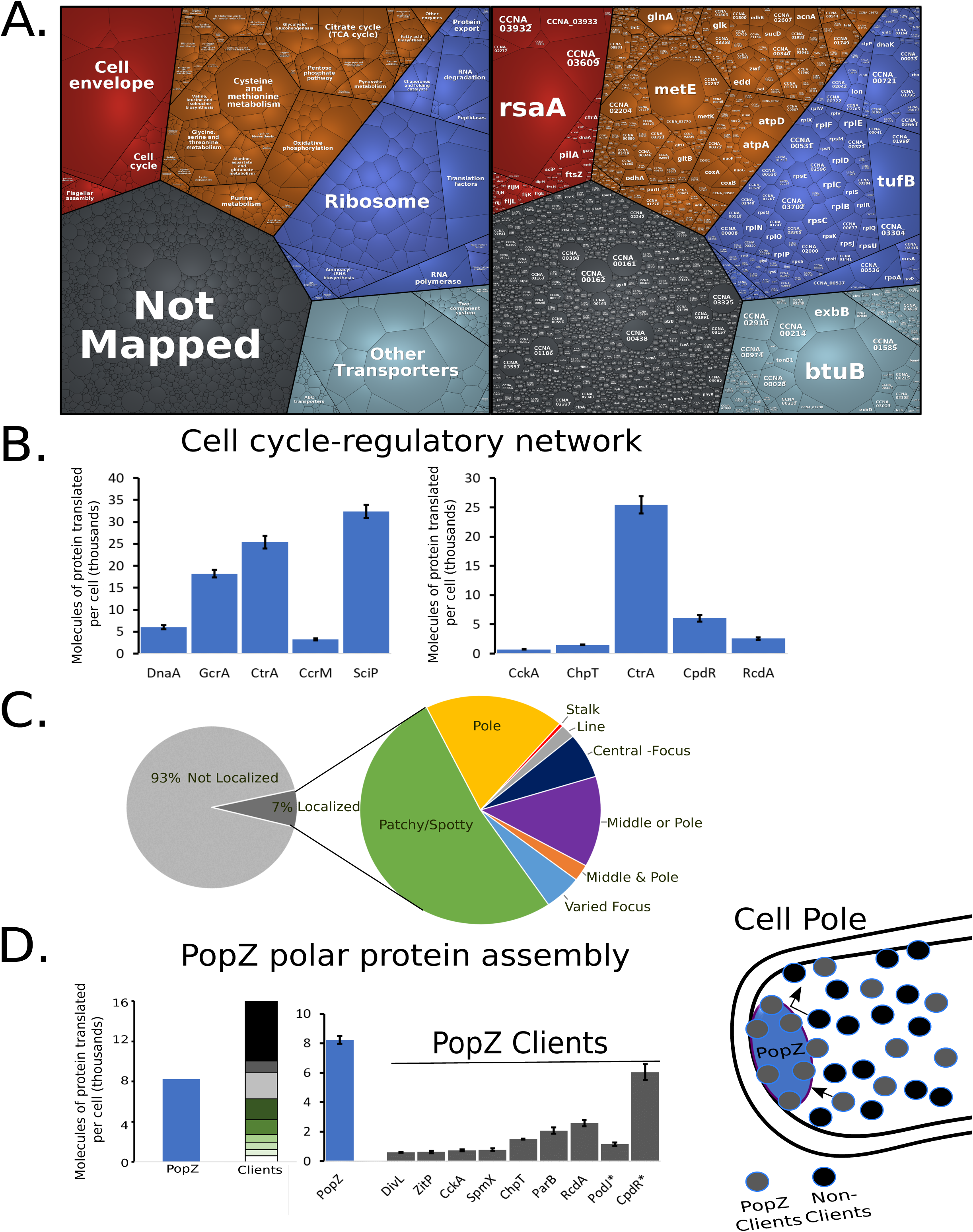
Global analysis of *C. crescentus* protein synthesis. A.) Proteomap with each polygon representing a single gene with area scaled to the fraction of ribosome protected mRNA footprints measured. Red is cellular processes, orange is metabolism, blue is genetic information processing, and light blue is environmental information processing, and grey are genes of unknown function. B.) Molecules of protein translated per cell for the cell cycle master-regulators (left) and CtrA regulatory network (right). C.) Left, fraction of ribosome protected mRNA footprints in localized (dark grey) or non-localized mRNAs (light grey). Right, zoomed in analysis of the fraction of proteins with different subcellular localization patterns based on (Werner et al., 2009). D.) Polar protein competition. Left, molecules of protein translated per cell for the polar protein scaffold PopZ and its known clients (Berge et al., 2016; Holmes et al., 2016; Zhao et al., 2018). Proteins with known proteolysis are highlighted with asterisks. Right, cartoon of polar protein assemblies with client proteins and non-client proteins indicated.

Interestingly, while the cell cycle is a major area of study in *C. crescentus*, the cell cycle genes only represent a small fraction of protein synthesis capacity (1.78%), yet these genes play a critical role in shaping the growth and division cycles. The cell cycle-regulatory circuit itself is composed of four transcription factors (DnaA, GcrA, CtrA, and SciP) and a DNA methylase (CcrM) whose spatiotemporal activation facilitate cell cycle progression (K. Lasker et al., 2016). For master-regulator proteins, the cell produces between ∼3,000-30,000 copies of each protein which corresponds to between 71-648 fold more proteins than the number of known DNA binding sites that they control (Zhou et al., 2015), with the exception of CcrM (Table S4). 3280 CcrM proteins are translated to methylate the 4542 GANTC sites per chromosome (Fig 2B). While the number of CcrM proteins are approximately 1/3 the number of GANTC sites present after DNA replication, CcrM is a processive enzyme (Berdis et al., 1998), suggesting that each CcrM may on average methylate ∼3 GANTC sites. While the number of CtrA proteins translated (25,400 proteins) corresponds closely with the amount measured in predivisional cells (18,000-22,000 (Judd, Ryan, Moerner, Shapiro, & McAdams, 2003; Spencer, Siam, Ouimet, Bastedo, & Marczynski, 2009)), CtrA is produced at a significantly higher level than its collection of regulatory kinases, phosphotransferases, and proteolytic adapters that control its cell cycle-dependent activity (Fig 2B). GcrA interacts with the RNA-polymerase/σ^70^ complex to activate transcription of target promoters (Haakonsen, Yuan, & Laub, 2015) where a ∼4-fold excess of GcrA is produced over σ^70^, suggesting that excess GcrA may accelerate binding to the RNA-polymerase holoenzyme (Wu et al., 2018) to facilitate subsequent recruitment of σ^70^.

As many proteins were found to have distinct subcellular patterns of protein accumulation in *C. crescentus* (Werner et al., 2009), we compared protein synthesis capacity to the localization patterns of proteins observed in this dataset (Fig 2C). 7% of protein synthesis occurs for “localized proteins” in *C. crescentus*. Of those localized proteins, most are “patchy/spotty”, while a significant fraction has a subcellular address where the protein accumulates (pole, stalk, or center) (Fig 2C). Many proteins are specifically required to form asymmetric polar protein complexes that function to determine cell fate upon division (K. Lasker et al., 2016). Many of these polarly localized proteins are recruited to the cell pole through the multimeric hub protein PopZ (Berge et al., 2016; Holmes et al., 2016; Zhao et al., 2018). Interestingly, by examining PopZ and its known client proteins, we find that PopZ is made in limiting amounts (Fig 2D), suggesting that the clients compete for PopZ binding *in vivo*.

### Analysis of vitamin B_12_ and methionine metabolism

Analysis of the most highly translated proteins found that RsaA, the surface layer protein, was the most highly translated protein in the cell (Fig 3A) (Lau, Nomellini, & Smit, 2010). Elongation factor Tu was the 3^rd^ most abundant cytoplasmic protein owing to its requirement to deliver aminoacyl-tRNAs to the ribosome during translation (Krab & Parmeggiani, 1998). Surprisingly, we also observed that the homolog of the B_12_ importer (*btuB* 2^nd^ highest) and methionine biosynthetic gene (*metE* 4^th^ highest) were among the most highly translated proteins. Vitamin B_12_ is an important enzymatic cofactor that in *C. crescentus* is used for the biosynthesis of methionine, dNTP production, tRNA modification, and isomerization of methylmalonyl-CoA to succinyl-CoA (Fig 3) (Zhang, Rodionov, Gelfand, & Gladyshev, 2009)*. C. crescentus* cannot synthesize B_12_ *de novo* but can import it through the BtuB protein (Menikpurage, Barraza, Melendez, Strebe, & Mera, 2019). In the cytoplasm, both MetE and MetH perform the rate-limiting step of methionine biosynthesis, where MetH requires B_12_ but has a higher specific activity than MetE (Fig 3B) (Frasca, Banerjee, Dunham, Sands, & Matthews, 1988; Whitfield, Steers, & Weissbach, 1970). Of note, both BtuB and MetE are translated at much higher levels than the other components related to methionine biosynthesis (Fig 3B). Both the *btuB* and *metE* genes are the only two genes in *C. crescentus* encoded with B_12_ riboswitches in their 5’ UTRs (Schrader et al., 2014). We tested the function of these riboswitches by creating 5’ UTR fusions to the mCherry gene driven by the vanillate promoter and subjecting the cells to various concentrations of B_12_ in the form of cyanocobalamin (Fig 3C, Table S5). Both the MetE and BtuB 5’ UTR reporters showed high translation in the absence of B_12_ and exhibited a B_12_ concentration dependent translational shutoff (Fig 3C). The *metE* riboswitch appears to be more sensitive to B_12_ concentration, with K_1/2_=0.062nM, while the *btuB* riboswitch K_1/2_=0.19nM, both in line with the concentrations found in aquatic ecosystems (Fig S3) (Benoit, 1957; Pommel, 1975). Taken together, these data show that *C. crescentus* cells are investing a large amount of their protein synthesis capacity towards B_12_ uptake and the B_12_ independent methionine pathway in the absence of B_12_. We therefore hypothesized that the cells are wasting energy in the absence of B_12_ by producing these very costly proteins.

**Figure 3.**
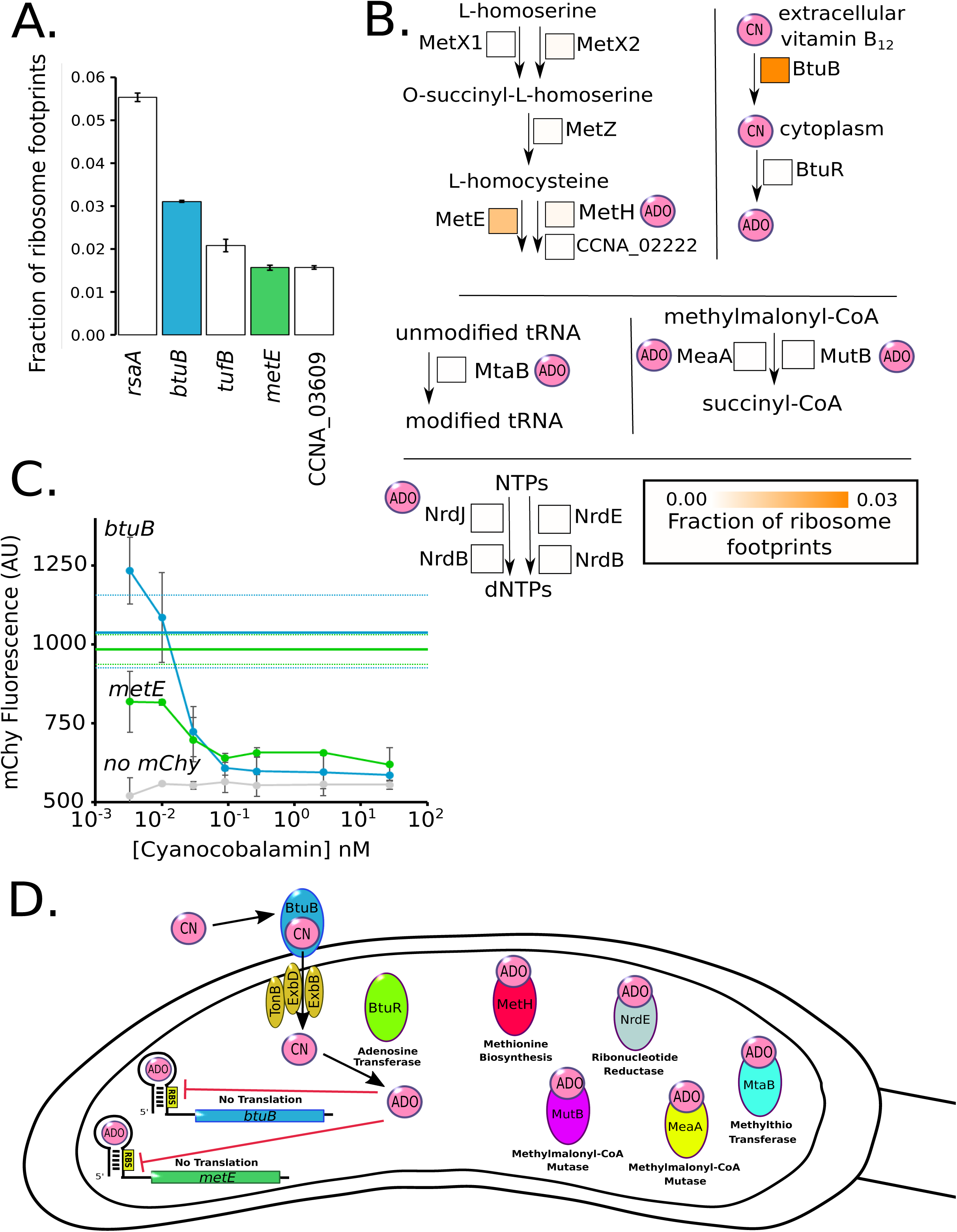
*C. crescentus* cells are starved for B_12_ in laboratory growth media. A.) Fraction of ribosome footprints for the most highly translated mRNAs in M2G. B_12_ related genes are colored. B.) Pathway of methionine biosynthesis, MetX1 (CCNA_03309), MetX2 (CCNA_00559), MetZ, (CCNA_02321), MetE (CCNA_00515), MetH (CCNA_02221), and CCNA_02222. Pathway for B_12_ utilization, BtuB (CCNA_01826), BtuR (CCNA_02321). Pathway for tRNA modification, MtaB, (CCNA_03798). Pathway for nucleotide reduction, NrdJ (CCNA_01966), NrdE (CCNA_03607), and NrdB (CCNA_00261). Pathway for succinyl-CoA biosynthesis MeaA (CCNA_03177), and MutB (CCNA_02459). Square boxes next to each enzyme contain an orange heat map which represents the fraction of ribosome footprints. ADO (adenosyl) and CN (cyano) refer to the upper ligand. C.) Negative regulation by B_12_ riboswitches on the *btuB* and *metE* genes. Translation reporters for the *btuB* and *metE* genes fused to mCherry assayed in M2G with the indicated concentrations of B_12_. Error bars represent standard deviation of mCherry fluorescence in three biological replicates of the B_12_ dilution series (n=3). Solid blue and green horizontal lines indicate the mChy fluorescence without vitamin B_12_ for *btuB* and *metE* respectively, and dashed lines indicate the standard deviation. D.) Cartoon of B_12_ regulated pathways in *C. crescentus*. *btuB* and *metE* genes contain negative regulatory B_12_ riboswitches. BtuB and BtuR are part of the B_12_ import and utilization pathway. MetH, NrdE, MeaA, MutB, and MtaB are B_12_ dependent enzymes for methionine biosynthesis.

To assess if the cells are wasting energy from the B_12_ related pathways, we examined the fitness of *C. crescentus* cells with disruptions in the non-essential components of these pathways in the absence of B_12_ (Fig 4A). For this we used Tn-seq datasets (Price et al., 2018) and analyzed B_12_ related genes whose disruption would not have polarity effects (single genes or last genes in operons) in M2G minimal media or PYE rich media, neither of which contain B_12_ in them (Poindexter, 1964; Schrader & Shapiro, 2015). In M2G minimal media, cells require the *metE* gene to make methionine (Menikpurage et al., 2019), while the other non-essential B_12_-related components showed increases in fitness when disrupted that were proportional to their protein synthesis costs (Fig 4A). In PYE rich media, which contains methionine in the peptone, we find that the *metE* gene is no longer essential, but instead its disruption leads to higher fitness. Indeed, all the components of the methionine pathway led to increases in fitness proportional to their protein synthesis cost when disrupted in PYE (Fig 4A). These data show that in the absence of B_12_, the excessive translation of these proteins leads to unnecessary costs of protein synthesis that limits the fitness of *C. crescentus* cells.

**Figure 4.**
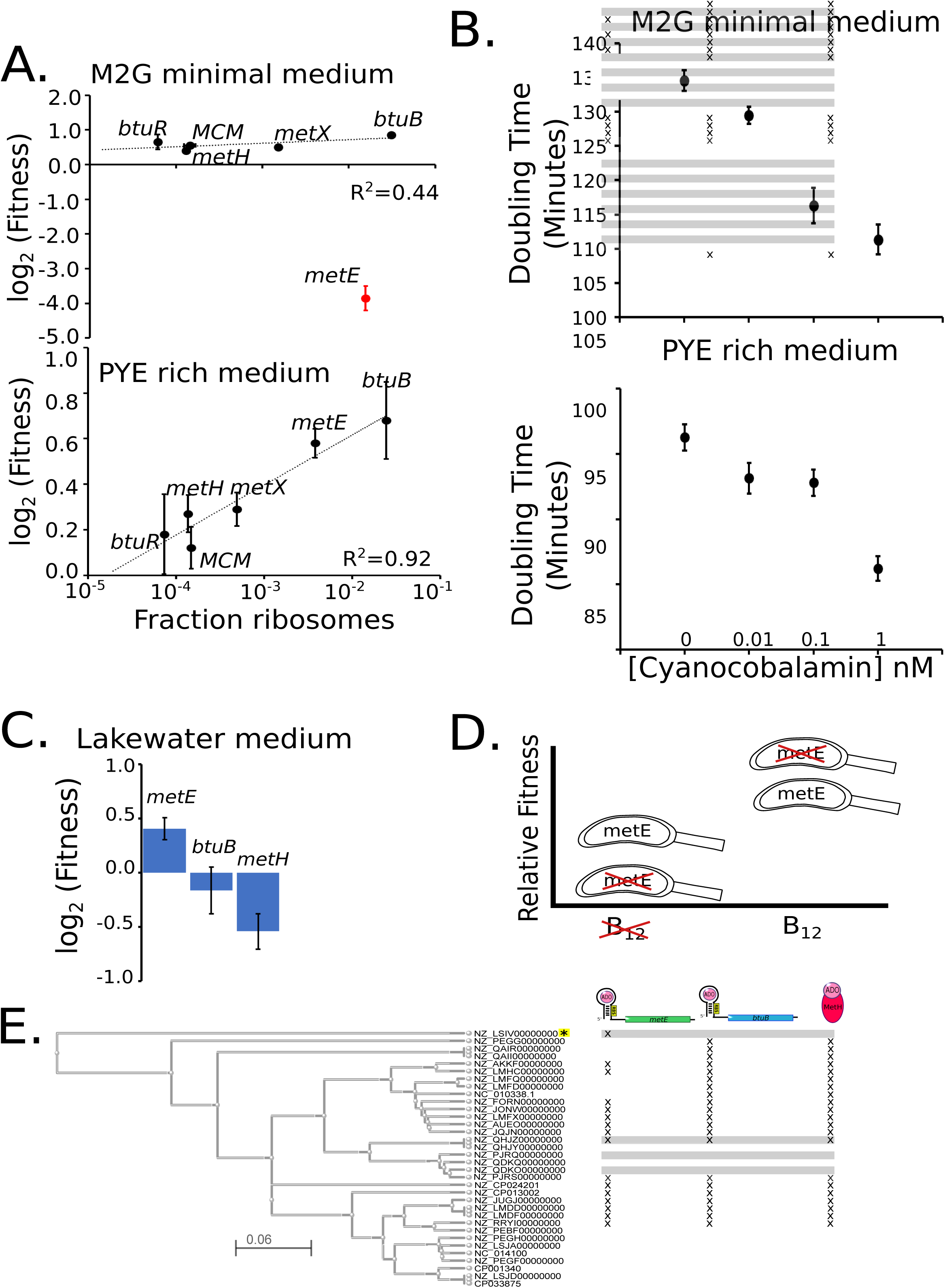
Excess protein synthesis rates for methionine biosynthetic genes correlates with fitness cost. A.) Protein synthesis cost measured in the fraction of ribosomes (Table 1) on the X-axis vs the Tn-seq derived fitness values for the for the *btuB* B_12_ importer and methionine biosynthetic genes under growth in minimal or rich media (Price et al., 2018) (biological replicates, n=2 for M2G, n=10 for PYE.). Black points represent non-essential genes for methionine biosynthesis and red points represent genes required for methionine biosynthesis under the specified growth condition. Error bars represent standard deviation. Curve fits are performed only on non-essential genes. B.) Doubling times of *C. crescentus* cells in M2G and PYE media with indicated concentrations of B_12_. Error bars represent the standard deviation of doubling time measurements (biological replicates, n=3 for each condition). C.) Tn-seq derived fitness values for *metE, btuB, and metH* under growth in Lake Michigan Lake water (Hentchel et al., 2019). D.) Fitness tradeoff of facultative vs obligate B_12_ scavenging. Relative fitness shown for species with *metE* (facultative) or without *metE* (obligate) in environments lacking or containing sufficient B_12_. E.) Phylogenetic tree of all *Caulobacter* species with completed genomes based on *btuB* and *metH* protein sequences. Each species is labeled by its NCBI Accession Identifier, and the scale represents the Kimura distance. Marks next to species represent the presence of a *metE, btuB*, or *metH* gene. All species have a predicted B_12_ riboswitch upstream of *btuB* and *metE* genes (not shown) except the species noted with the yellow asterisk.

To test the effects of B_12_ on *C. crescentus* cell growth we examined the growth rate of cells cultured in M2G media containing B_12_ in the form of cyanocobalamin. Here we observe faster growth in a B_12_ concentration dependent manner (Fig 4B, Table S6) with up to a 21% faster doubling time observed at 1 nM B_12_ in M2G media. 11% acceleration of cell growth rates also occurs in PYE media, which contains methionine in the peptone, suggesting that the growth enhancement of B_12_ is likely caused in part from reduced protein synthesis costs from *btuB* and *metE* riboswitches. We attempted to separate B_12_’s effects on methionine synthesis from its effects on other pathways by addition of exogenous methionine in the presence or absence of B_12_, but exogenous methionine leads to a dramatic decrease in growth by an unknown mechanism (Ferber & Ely, 1982). Overall, we find that B_12_ significantly enhances the growth rate of NA1000 cells.

## Discussion

### Absolute quantitation of protein synthesis in *C. crescentus*

As the cost of protein synthesis is a significant output of the cell’s energy, absolute quantitation of translation is a powerful method to measure gene expression and resource allocation. Here we present an absolute protein synthesis resource for *C. crescentus* generated by ribosome profiling which will be vital for systems modeling efforts of the *C. crescentus* cell cycle (Lin, Crosson, & Scherer, 2010; Murray, Panis, Fumeaux, Viollier, & Howard, 2013; Shen et al., 2008; Subramanian & Tyson, 2017) and in the subsequent optimization of synthetic *Caulobacter* genomes (Christen, Deutsch, & Christen, 2015).

Across the proteome the mRNA translation resource allocation showed 1.4% of the translation machinery is dedicated to translation of cell cycle-regulatory genes. We observed that for all the cell cycle-master regulators, except for CcrM, that the number of proteins translated dramatically exceeds the total number of DNA binding sites in the genome (Fig 2B). We hypothesize that the high concentrations of these factors may facilitate rapid activation of target gene transcription during each phase of the cell cycle.

Approximately 7% of protein synthesis capacity is dedicated to genes whose protein products were found to be localized in one of several modes of subcellular organization (Fig 2C) (Werner et al., 2009). Recent reports suggest that some of these foci are formed by liquid-liquid phase separation of the proteins into membraneless organelles (Al-Husini, Tomares, Bitar, Childers, & Schrader, 2018; Banani, Lee, Hyman, & Rosen, 2017). For *C. crescentus* BR-bodies, the concentration of the condensate forming protein RNase E (6.3 μM) appears to correspond closely to the transition boundary for liquid-liquid phase separation (Al-Husini et al., 2018), potentially allowing control of the assembly of these bodies. Interestingly, we observed that the polar protein scaffold PopZ, which facilitates recruitment of asymmetrically localized signaling proteins to the cell poles (Berge & Viollier, 2018; Holmes et al., 2016; Zhao et al., 2018), is present at approximately 1/2 the concentration of its client proteins suggesting that clients compete for PopZ access (Fig 2D). Dynamic competition of clients for PopZ may be important for the ordered assembly of unique proteins at each cell pole and may impact the spatial activation of downstream signaling outputs (Childers et al., 2014; Holmes et al., 2016; Keren Lasker et al., 2018).

### Implications of B_12_ scavenging pathway on environmental fitness

B_12_ is an important enzymatic cofactor that is required for the activity of enzymes involved in biosynthesis of methionine, dNTP production, tRNA modification, and isomerization of methylmalonyl-CoA to succinyl-CoA (Fig 3B,D) (Zhang et al., 2009). Like many bacteria, *C. crescentus* cannot produce B_12_ but can scavenge it form the environment (Menikpurage et al., 2019), where it can increase the growth rate of *C. crescentus* by up to 21% (Fig 4B). b*tuB*, the B_12_ importer, and *metE*, the B_12_-independent methionine synthase, are among the most highly expressed genes accounting for ∼4.5% of all protein synthesis capacity (Fig 3A). To counteract the protein synthesis demand, *btuB* and *metE* genes also contain B_12_ riboswitches that reduce translation when sufficient B_12_ enters the cytoplasm (Fig 3C). Freshwater bodies typically have B_12_ concentrations in the range from 0.11 nM to below the level of detection (<0.1 pM) (Benoit, 1957; Cavari & Grossowicz, 1977; Daisley, 1969; Kouichi Ohwada, 1973; Kouichi Ohwada & Taga, 1972; Pommel, 1975) suggesting that high levels of BtuB may help facilitate import. Importantly, the conserved *metE* and *btuB* riboswitches are sensitive to B_12_ in physiologically relevant ranges (K_1/2_=0.062 nM, 0.19 nM, respectively) and their different sensitivities suggest that *metE* translation would be downregulated before shutting off the *btuB* importer. At high concentrations of B_12_ that saturate the riboswitches (Fig 4B), a significant portion of the increase in growth rate is likely due to the liberation of protein synthesis resources on these two highly expressed genes (Fig 3A,C). When grown in M2G or PYE media lacking B_12_, disrupting the *btuB* gene increases fitness by freeing up wasted protein synthesis resources (Fig 4A). Similarly, *metE* disruption leads to increased fitness in PYE which contains methionine, but *metE* disruption becomes essential in M2G minimal media as its required to make methionine (Fig 4A). Why then does *C. crescentus* have a facultative B_12_ lifestyle containing both B_12_ dependent and independent methionine biosynthesis pathways?

Perhaps B_12_ independent and B_12_ dependent pathways exist to buffer fluctuations in environmental B_12_ concentrations. The concentration of available B_12_ in a freshwater body shows variation of up to 40 fold between different sampling locations and at the same sampling location at different times (Cavari & Grossowicz, 1977; Daisley, 1969; Kouichi Ohwada, 1973; K Ohwada, Otsuhata, & Taga, 1972; Kouichi Ohwada & Taga, 1972; Pommel, 1975). Having both methionine biosynthesis pathways adds flexibility to generate methionine in either high or low B_12_ conditions, however, these pathways have different protein synthesis costs. The B_12_ independent pathway requires 1.57% of the total protein synthesis capacity to make sufficient MetE, while the the B_12_-dependent MetH pathway uses only 0.156% (Fig 3B). Interestingly, disrupting the *metE* gene in lakewater leads to a fitness advantage, however, disrupting *btuB* or *metH* leads to a fitness decrease (Fig 4C) (Hentchel et al., 2019). Although not measured directly, the gene fitness signature from the experiment leads us to infer that physiologically relevant B_12_ concentrations were present in the sampled lake water. The increased fitness of *metE* disruptions in lakewater suggest that increased biosynthetic flexibility comes with a negative fitness cost from wasted protein synthesis resources on MetE (Fig 4D). 47% of fully sequenced *Caulobacter* species have lost the *metE* gene but not the *btuB* and *metH* genes, suggesting that the observed environmental fluctuations in B_12_ concentration (Daisley, 1969; Pommel, 1975) alter the selective pressure on *metE* (Fig 4E).

Surprisingly, a recent survey of available metagenomic 16S rRNA sequencing data showed that *Caulobacter* are more abundant in soil/compost than in aquatic ecosystems (Wilhelm, 2018). Soil has been shown to have B_12_ levels that can range from 20 nM to 0.3 nM, correlated with levels of organic matter (Duda, Malinska, & Pedziwilk, 1957), while bodies of freshwater typically have B_12_ concentrations in the range 0.11 nM to below the level of detection (0.1 pM) (Benoit, 1957; Cavari & Grossowicz, 1977; Kouichi Ohwada, 1973; K Ohwada et al., 1972; Kouichi Ohwada & Taga, 1972; Pommel, 1975). The higher B_12_ concentration in soil will enhance the growth rate and may explain the increase in relative abundance in this environment.

## Materials and Methods

### Bacterial strains and cell growth

A list of all bacterial strain used here can be found in table S7. *C. crescentus* cells were grown in M2G or PYE growth media (Schrader & Shapiro, 2015) and supplemented with the appropriate antibiotic concentrations (Thanbichler, Iniesta, & Shapiro, 2007). *E. coli* cells used for cloning were grown in LB media and supplemented with the appropriate antibiotics.

### Ribosome profiling

Ribosome profiling was performed similar to (Schrader et al., 2016; Schrader et al., 2014) except contaminating rRNA fragments generated during MNase digestion were depleted to allow deeper quantitation of resulting mRNA translation similar to (Li et al., 2014). For a detailed protocol of the procedure see (Aretakis et al., 2018). 500mL of NA1000 cells were grown in M2G media to an OD600 of 0.5 and treated with 100μg/mL Chloramphenicol for 2 min, then harvested by centrifugation and flash frozen in liquid nitrogen. Cells were then lysed on a mixer mill (Retsch mm400) at 6 cycles of 3 min at 15Hz, thawed, membranes pelleted, and the supernatant was footprinted by addition of MNase (Roche). After footprinting, MNase was quenched with EGTA, and samples were separated by sucrose gradient fractionation. 70S peaks were purified, phenol chloroform extracted, and ethanol precipitated (Aretakis et al., 2018). Resulting mRNA fragments were size selected by 10% acrylamide 1XTBE/7M Urea PAGE, end repaired, 3’ adapter ligated, reverse transcribed, circularized, and depleted of rRNA fragments (Aretakis et al., 2018; Ingolia et al., 2009; Li et al., 2014). Ribosomal RNA cDNA fragments were removed using biotin-linked DNA oligos (oCaulo1: 5’/5Biosg/CGCTTACGGGGCTATCACCCA, oCaulo2: 5’/5Biosg/TGGCAACTAATCACGAGGGTT, oCaulo3: 5’/5Biosg/CTCATCTGGTTGCCCAAAAGA, oCaulo4: 5’/5Biosg/TGGTTCAGGAATATTCACCTG) and MyOne Streptavidin C1 Dynabeads (Invitrogen) as in (Li et al., 2014). Resulting circular cDNAs were amplified by PCR using Phusion DNA polymerase (Fermentas) with indexing primers (Ingolia, Brar, Rouskin, McGeachy, & Weissman, 2012), pooled together, and sequenced on an Illumina hiseq 2000. Data for three ribosome profiling replicates were deposited in the gene expression omnibus under accession number GSE126485. The three M2G replicates were further analyzed together with a PYE dataset from (Schrader et al., 2014).

Ribosome footprint reads were mapped to the genome as center weighted reads(Oh et al., 2011), and extreme fast and slow codons were corrected for by winsorization of the bottom 5% and top 95% of nucleotides. The resulting fraction of ribosome footprints (ϕ_*i*_of each gene (*i*) compared to the total ribosome footprint total were calculated and converted into the number of molecules protein translated per cell (*ki*) by 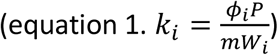 where P is the average protein mass per cell and *mW*_*i*_ is the protein product’s molecular weight as originally described in (Li et al., 2014). Average protein mass per cell (P) was measured as follows: 5mL mid log cultures of NA1000 cells were grown in M2G or PYE media overnight in a roller wheel at 28°C. Once the cells reached an 0D600nm of 0.3, a 100μL aliquot of the cells were diluted and counted on PYE plates to measure the number of viable cells and another aliquot was saved for protein concentration measurements. To measure protein content, 100μL cells were spun down in a microcentrifuge at 14,000rpm for 30 seconds, the supernatant removed, and the remaining cell pellets were resuspended in 100μL of 1X Laemmlli sample buffer lacking any dyes. After resuspension, samples were boiled for 5 min at 95°C then placed on ice. Lysate protein concentrations were measured using the Pierce™ 660nm Protein Assay (Thermo Fisher) and comparing the lysate A600 with a linear curve fit of BSA standards. The mass of protein from the sample was then divided by the number of viable cells to yield the average protein mass/cell. The resulting average protein mass per cell were: 492×10^−15^g/cell ± 170 in M2G and 519 x10^−15^g/cell ± 228 in PYE.

### Absolute quantitation of C-terminal YFP fusions

5mL mid log cultures of NA1000 cells harboring C-terminal eYFP fusions (gift of the Shapiro Lab, Stanford U.) were grown in M2G or PYE media overnight in a roller wheel at 28°C. Once the cells reached an 0D600nm of 0.3, cells were spotted on M2G agarose pads for imaging. Images were collected on Leica DM6000B microscope with a Hamamatsu C9100 EM-CCD camera and a 100X PH3 PlanApo 1.40 NA objective with in a Semrock model 2427A YFP filter cube with 100ms exposure time. Fluorescence intensity was quantified using ImageJ by segmenting the cells and measuring the average pixel intensity of the cell area. Background intensity was subtracted using the NA1000 average YFP pixel intensity. For each fusion strain, a minimum of 50 cells were used for the analysis with a minimum of two technical replicates. As MipZ molecules per cell had been previously measured (Thanbichler & Shapiro, 2006) by quantitative western blot, we converted the MipZ-YFP pixel intensity to the number molecules/cell and multiplied this conversion factor by the YFP intensities of all other C-terminal YFP fusions. The average across replicates and the standard deviation (σ) are reported in table S2.

### Doubling time measurements

Treatments were started from log phase cultures grown overnight in the absence of cyanocobalamin (Sigma-Aldrich) and diluted in fresh media to an OD_600_ of 0.05. Each treatment was split into a separate flask and the correct amount of cyanocobalamin was added to the concentrations of 1 nm, 0.1 nm, 0.01 nm and 0 nm. 50 mL from each treatment was added to three different 250 mL Kimex flasks, for three replicates of each of the four treatments. An initial OD_600_ measurement was taken of each replicate using a cuvette and a nanodrop spectrophotometer. The 12 – 250 mL flasks were then placed in a 28 °C shaker incubator at 250 RPM. OD_600_ time points of each flask were taken throughout the logarithmic growth phase. An exponential regression of the log phase time points was used to calculate the doubling time of each replicate.

### Translation reporter assay

JS417, JS423, and JS440 strains were started from log phase cultures grown overnight in the absence of cyanocobalamin and diluted in media with vanillate and antibiotic to an OD_600_ of 0.05. A dilution series of each strain was used to fill tubes with 2 mL of culture at each cyanocobalamin concentration; 27 nm, 2.7 nm, 0.9 nm, 0.3 nm, 0.1 nm, 0.033 nm, and 0 nm. The 21 2-mL-cultures were then grown and induced over an eight-hour period by placing the tubes in a 28 °C shaker incubator at 250 RPM for 8 hours. After eight-hours, 2 μL from each culture was pipetted onto an M2G-agrose pad on a microscope slide. Each treatment was imaged on a microscope using both phase contrast and a mChy filter cube. Average fluorescent intensities were calculated using MicrobeJ (Ducret, Quardokus, & Brun, 2016) across a minimum of 100 cells.

### B_12_ homolog identification and phylogenetic tree mapping

Protein sequences for *btuB, metH,* and *metE* were determined for each *Caulobacter* species with a complete genome by using the NCBI Basic Local Alignment Search Tool (BLAST) with default settings by searching the protein sequence of each NA1000 gene for homologs with an E-score of <10^−19^ (Sayers et al., 2019). The *btuB* and *metH* genes were then used to generate the phylogenetic tree using the NCBI Genome Workbench software and the MUSCLE multiple sequence alignment package (Edgar, 2004). The tree is a maximum likelihood generated with the default settings from MUSCLE. Riboswitches were identified using rfam (Gardner et al., 2011) and searching the upstream regions of *btuB* and *metE* genes (up to 1000bp upstream of their predicted operons).

### Proteomap generation

Categories were taken from predefined Kyoto Encyclopedia of Genes and Genomes (KEGG) categories (Kanehisa & Goto, 2000). Categories were then ranked based on priority, and any gene that may be present in more than one category were deleted from those with lower priority. The 200 most numerous proteins were then hand checked. Any that had not be automatically assigned to a KEGG category were compared against other organisms to place them in their most appropriate category. The categorized genes along with the ribosome profiling data was used to create the Proteomap (Liebermeister et al., 2014).

## Acknowledgements

The authors thank: Peter Chien for sharing absolute protein concentrations of DnaA, Lon, and CcrM, Adam Perez for sharing the tipN-YFP strain, Paola Mera for sharing data on *btuB* and for thoughtful discussion, and members of the Higgs lab for thoughtful discussions. This work was supported by NIH R35 GM124733 to JMS, WSU start-up funds to JMS, and a Chemical Biology Interface research experience award WSU to JRA.

