## Supplemental Information for "Absolute measurements of mRNA translation in *C. crescentus* reveal important fitness costs of vitamin B_12_ scavenging"

### Supplementary Figures

#### Figure S1

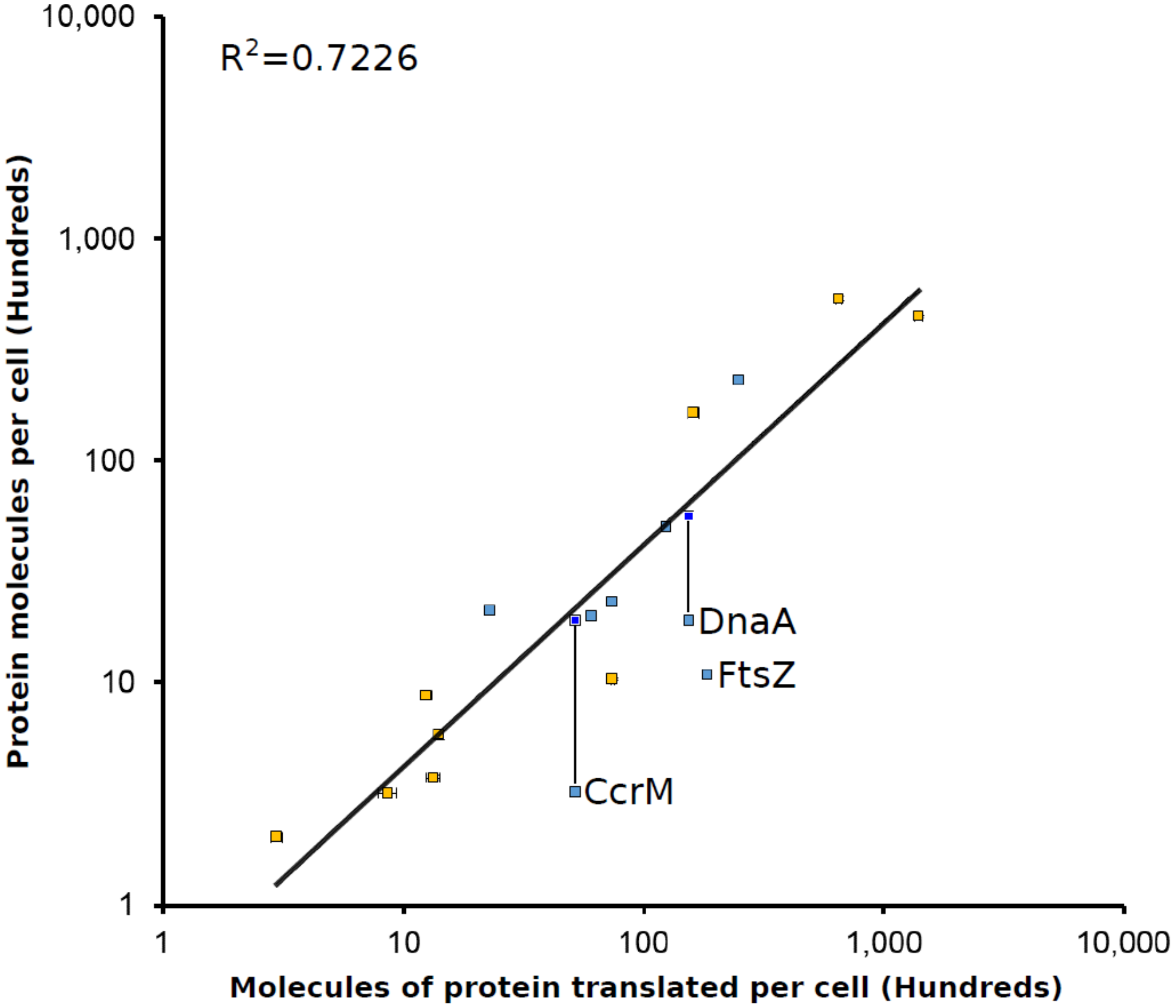


Fig S1. Absolute protein levels of unsynchronized cells in PYE media measured by western blot (blue) or YFP fusions (orange) compared to the absolute molecules of protein translated per cell calculated by ribosome profiling. Vertical error bars indicate the standard deviation in YFP intensity or standard deviation for the western blots. FtsZ, CcrM, and DnaA are indicated as their protein levels are under proteolytic control [14]. CcrM and DnaA protein levels in a strain lacking their protease, Lon, are indicated in dark blue (Peter Chien personal communication). Data in Tables S1 and S2.

**Figure S2**

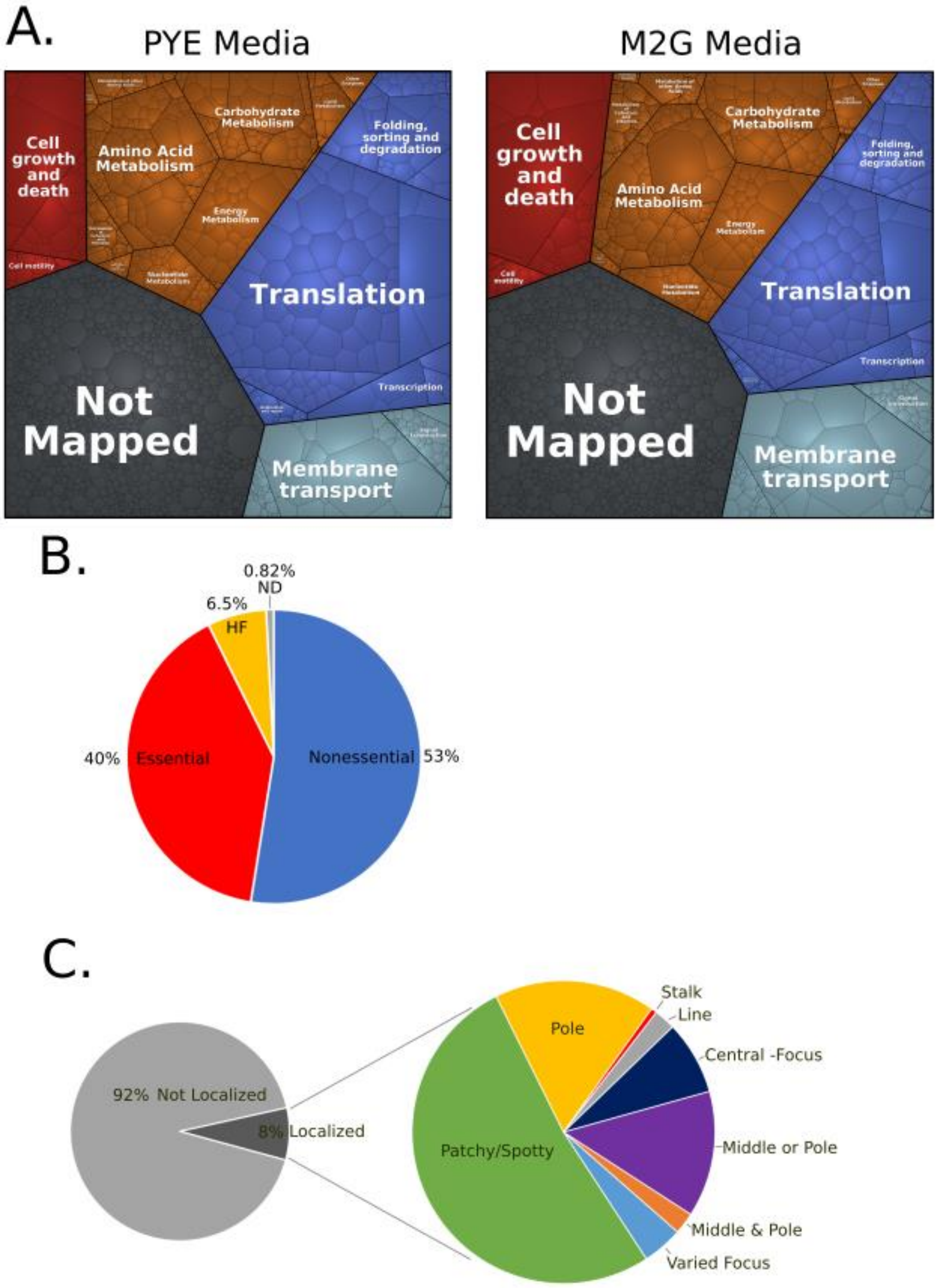
A.
PYE Media
Cell growth and death
Amino Acid Metabolism
Carbohydrate Metabolism
Folding, sorting and degradation
Energy Metabolism
Cell motility
Nucleotide Metabolism
Translation
Transcription
Not Mapped
Membrane transport
M2G Media
Cell growth and death
Amino Acid Metabolism
Carbohydrate Metabolism
Folding, sorting and degradation
Energy Metabolism
Cell motility
Nucleotide Metabolism
Translation
Transcription
Not Mapped
Membrane transport
B.
0.82% ND
6.5% HF
40% Essential
Nonessential 53%
C.
92% Not Localized
8% Localized
Pole
Stalk
Line
Central -Focus
Middle or Pole
Middle & Pole
Varied Focus
Patchy/Spotty

Fig S2. PYE fractional protein synthesis Proteomap.

Global analysis of C. crescentus protein synthesis in PYE and M2G media. A.) Left, Proteomap of cells grown in PYE media with the area scaled to the fraction of ribosome protected mRNA footprints measured. Right, Proteomap of cells grown in M2G media shown at the same level as cells grown in PYE Media with KEGG categories (Kanehisa & Goto, 2000) B.) Fraction of ribosome protected mRNA footprints by essentiality. Red is essential genes, blue is nonessential, yellow is high fitness genes, and grey is genes that were not determined (Christen et al., 2011). C.) Left, fraction of ribosome protected mRNA footprints in localized (dark grey) or non-localized mRNAs (light grey) for cells grown in PYE media. Right, zoomed in analysis of the fraction of proteins with different subcellular localization patterns based on (Werner et al., 2009).

**Figure S3**

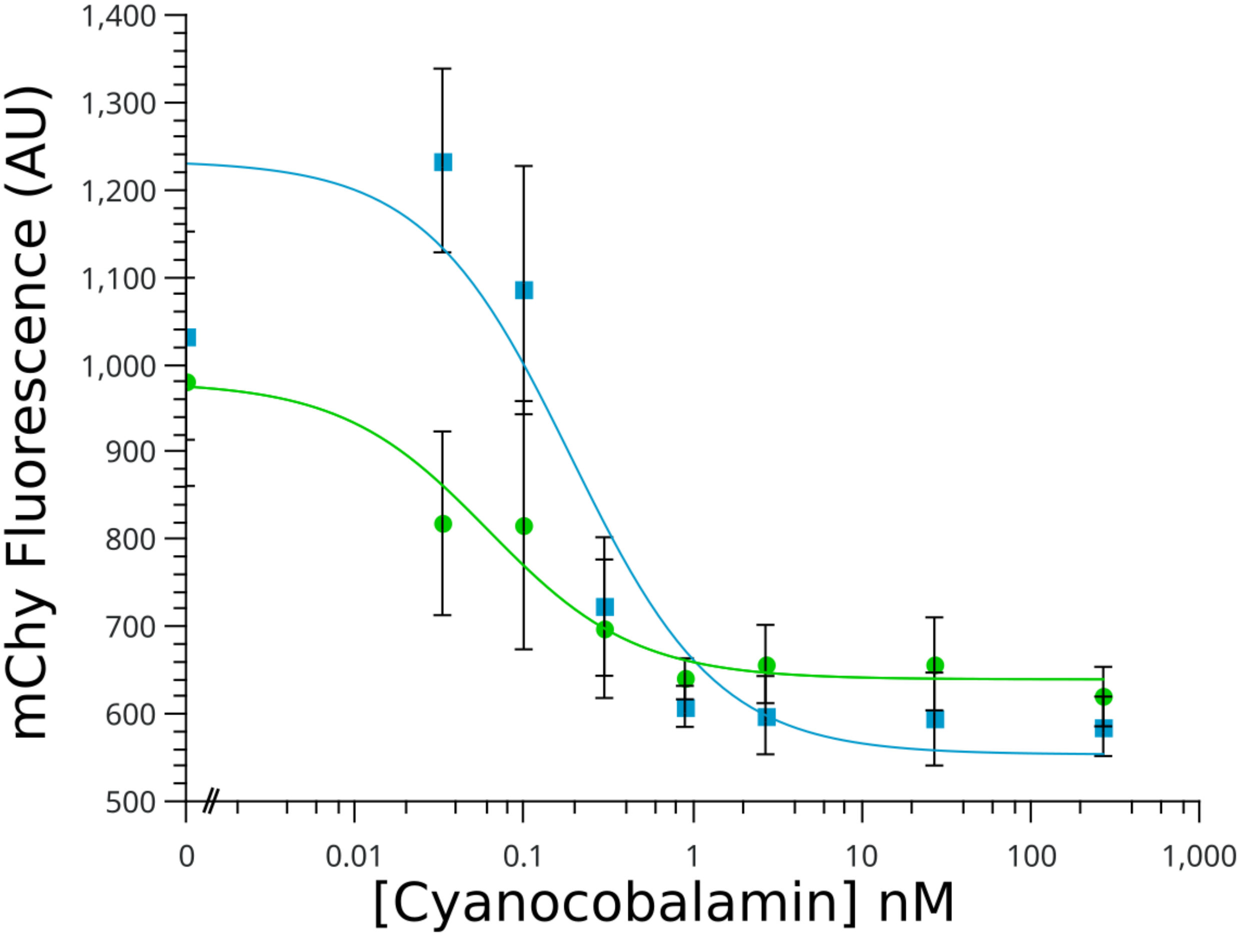


Fig S3. Non-linear curve fit for $B_{12}$ dependent riboswitch repression. mCherry intensities at each cyanocobalamin concentration were fit to a michaelis-menton equation using QTIplot software to determine the $K_{1/2}$ of cyanocobalamin. $R^2$ values were 0.95 for *metE* (green) and 0.85 for *btuB* (blue) respectively.

### Supplementary Tables

Table S1. Ribosome profiling values (see attached excel file).

Table S2. Average molecules of protein translated per cell and average proteins per cell.

| **M2G** | Ribosome profiling | | YFP-intensity | |
|---|---|---|---|---|
| Protein | Average molecules of protein translated per cell | σ | Average Proteins per cell | σ |
| B'-YFP | 1.38E+04 | 8.17E+02 | 1.64E+04 | 4.37E+02 |
| CckA-YFP | 7.34E+02 | 6.72E+01 | 6.37E+02 | 1.15E+02 |
| DnaB-YFP | 4.33E+02 | 6.61E+01 | 4.56E+02 | 1.57E+01 |
| HolC-YFP | 4.16E+02 | 7.74E+01 | 3.86E+02 | 9.44E+01 |
| Hu2-YFP | 5.58E+04 | 5.85E+03 | 4.20E+04 | 1.98E+03 |
| L1-YFP | 2.99E+04 | 1.90E+03 | 4.65E+04 | 1.95E+03 |
| MipZ-YFP | 2.41E+03 | 1.13E+02 | 1.03E+03 | 1.91E+02 |
| SMC-YFP | 2.50E+02 | 1.63E+01 | 2.21E+02 | 9.85E+01 |
| TipN-YFP | 9.70E+02 | 9.20E+01 | 6.23E+02 | 4.80E+01 |
| **PYE** | Ribosome profiling* | | YFP-intensity | |
| Protein | Average molecules of protein translated per cell | σ | Average Proteins per cell | σ |
| L1-YFP | 5.36E+04 | ND | 4.65E+04 | 1.39E+03 |
| B'-YFP | 1.64E+04 | ND | 1.64E+04 | 8.22E+01 |
| Ccka-YFP | 8.77E+02 | ND | 6.37E+02 | 1.11E+01 |
| DnaB-YFP | 5.79E+02 | ND | 4.56E+02 | 3.62E+01 |
| HolC-YFP | 3.17E+02 | ND | 3.86E+02 | 1.62E+01 |
| Hu2-YFP | 4.45E+04 | ND | 4.20E+04 | 9.78E+02 |
| Mipz-YFP | 1.03E+03 | ND | 1.03E+03 | 1.33E+01 |
| Smc-YFP | 2.01E+02 | ND | 2.21E+02 | 1.92E+00 |
| TipN-YFP | 3.72E+02 | ND | 6.23E+02 | 1.18E+01 |

*Schrader JM *et al*., PLoS Genetics (2014)

Table S3. Fraction of total protein synthesis for each KEGG category.

| Category | PYE | M2G |
|---|---|---|
| Cell growth and death | 6.45% | 11.26% |
| Cell motility | 1.19% | 1.09% |
| Membrane transport | 9.23% | 11.94% |
| Signal transduction | 1.38% | 1.30% |
| Folding, sorting and degradation | 5.88% | 5.08% |
| Replication and repair | 0.62% | 0.46% |
| Transcription | 2.62% | 2.37% |
| Translation | 22.82% | 15.88% |
| Amino Acid Metabolism | 8.11% | 8.54% |
| Carbohydrate Metabolism | 5.51% | 6.09% |
| Energy Metabolism | 4.54% | 4.47% |
| Glycan Biosynthesis and Metabolism | 0.27% | 0.24% |
| Lipid Metabolism | 1.03% | 0.93% |
| Metabolism of Cofactors and Vitamins | 1.09% | 1.53% |
| Metabolism of other Amino Acids | 0.90% | 0.97% |
| Metabolism of Terpenoids and Polyketides | 0.19% | 0.15% |
| Nucleotide Metabolism | 1.91% | 1.69% |
| Other Enzymes | 0.43% | 0.41% |
| Xenobiotics Biodegradation and Metabolism | 0.05% | 0.07% |
| Not Mapped | 25.77% | 25.52% |

Table S4. Comparison of absolute translation level to DNA binding sites.

| Gene | Molecules translated per cell | σ | DNA Sites in cell cycle-regulated promoters | Total DNA binding sites | References |
|---|---|---|---|---|---|
| *dnaA* | 6000 | 459 | 77 | 84 | (Taylor, Ouimet, Wargachuk, & Marczynski, 2011; Zhou et al., 2015) |
| *gcrA* | 18200 | 844 | 94 | 217 | (Fioravanti et al., 2013; Haakonsen, Yuan, & Laub, 2015; Zhou et al., 2015) |
| *ctrA* | 25400 | 1480 | 183 | 187 | (Fiebig et al., 2014; Zhou et al., 2015) |
| *ccrM* | 3280 | 233 | 96 | 4542 | (Kozdon et al., 2013; Zhou et al., 2015) |
| *sciP* | 32400 | 1480 | 61 | 61 | (Fumeaux et al., 2014; Zhou et al., 2015) |

Table S5. Fluorescent intensity measurements in arbitrary units collected from the translation reporters in M2G media with different cyanocobalamin concentrations.

| | Cyanocobalamin concentration (nM) | | | | | | | |
|---|---|---|---|---|---|---|---|---|
| | 0 | 0.033 | 0.10 | 0.30 | 0.90 | 2.7 | 27 | 270 |
| *btuB* | 951 | 1230 | 1080 | 723 | 609 | 599 | 595 | 586 |
| σ | 165 | 105 | 142 | 79.3 | 23.5 | 44.8 | 53.2 | 34.0 |
| *metE* | 980 | 818 | 817 | 698 | 640 | 657 | 657 | 620 |
| σ | 46.8 | 96.3 | 9.31 | 70.2 | 13.5 | 15.1 | 3.10 | 52.0 |
| MCS | 536 | 520 | 558 | 553 | 565 | 553 | 557 | 555 |
| σ | 22.2 | 57.3 | 5.77 | 11.8 | 34.0 | 34.8 | 35.9 | 14.1 |

Table S6. Doubling time measurements at different concentrations of cyanocobalamin.

| | Cyanocobalamin concentration (nM) | | | |
|---|---|---|---|---|
| | 0 | 0.01 | 0.1 | 1 |
| M2G (Minutes) | 135 | 129 | 116 | 111 |
| σ | 1.55 | 1.24 | 2.65 | 2.24 |

| | | | | |
|---|---|---|---|---|
| PYE (Minutes) | 101 | 98.1 | 97.8 | 91.2 |
| σ | 1.02 | 1.18 | 1.00 | 0.952 |

Table S7. List of bacterial strains.

| Strain | Genotype | Source |
|---|---|---|
| NA1000 | Synchronizable variant of CB15 | (Evinger and Agabian, 1977) |
| JS417 | NA1000 pRV(*btuB*_5'UTR)CHYC-2 Kan$^R$ | This Study |
| JS423 | NA1000 pRV(*metE*_5'UTR)CHYC-6 Chlor$^R$ | This Study |
| JS440 | NA1000 pRVMCS-2 Kan$^R$ | This Study |
| JS290 | NA1000 *L1::L1-yfp* Gent$^R$ | (Bayas et. al. 2018) |
| JS441 | NA1000 β'*::* β'*-yfp* Spec$^R$ Strep$^R$ | This Study |
| NJH429 | NA1000 *cckA::cckA-yfp* RifR | (Iniesta *et al*., 2010) |
| LS3587 | NA1000 dnaB*::dnaB-yfp* Kan$^R$ | (Rasmus *et al*., 2001) |
| LS3586 | NA1000 *holC::holC-yfp* Kan$^R$ | (Rasmus *et al*., 2001) |
| MS307 | NA1000 *hu2::hu2-yfp* Kan$^R$ | (Lee *et al*., 2011) |
| MT97 | NA1000 *mipZ::mipZ-yfp* | (Thanbichler and Shapiro, 2006) |
| LS3394 | NA1000 *smc::smc-yfp* Kan$^R$ | (Rasmus and Shapiro, 2003) |
| *tipN-YFP* | NA1000 *tipN::tipN-yfp* | (Gift from Adam Perez) |

#### Strain Construction

**JS417**

This strain was generated by purchasing *btuB*_5'UTR gBlock construct (IDT) for the +1 TSS site through the start codon of the *btuB* gene.

*btuB*_5'UTR gBlock:
AAGCGTTCAATTGGATCCAATCTTGACGTCCGTTTGATTACGATCAAGATTGGATCCAGCGTCAGGTTCCTCGAAA
GAGGATGAAAAGGGAACGAGGTTGAAGACCTCGGCTGCCCCCGCAACTGTAAGCGGCGAGCTTCGCGTCACATG
CCACTGGGCCCAAAAGGCCTGGGAAGGCGACGCCCAGAAGCATTGACCCGTGAGCCAGGAGACCTGCCCGGCGC
AGTCGTTCATCGCTCGGCCGGGGTGCGCCGAACGAACGGGATCTCCCGAGAAACGACAGTCAACAGGCCGCGCG
ACGGCCTGAGCGTCCGCGTCTTCGCGGGCGGTCGGGAGGTCGCGTGGGTCGTTCATAACGGGAAGACTGTATTAT
GTTAATTAATATGCATGGTAC

The plasmid pRVChyC-2 (Thanbichler, Iniesta, & Shapiro, 2007) was cut with MfeI and PacI, and the gblock segment was inserted into the plasmid by gibson assembly. Next, the resulting plasmid was transformed into E. coli DH5 alpha cells and selected on LB-kan plates. The resulting kanR colonies were then screened by PCR for the insert and verified by sanger sequencing (genewiz). The purified plasmid was then transformed into NA1000 cells by electroporation and plated on PYE-kan plates. The resulting colonies were screened for mChy fluorescence after induction with vanillate.

**JS423**

This strain was generated by purchasing *metE*_5’UTR gBlock construct (IDT) for the +1 TSS site through the start codon of the *metE*.

*metE*_5’UTR gBlock:
AAGCGTTCAATTGGATCCAATCTTGACGTCCGTTTGATTACGATCAAGATTGGATCCAGTCGTGGTCTGCGGACGT
TCGCGTCCGGAGCTAAGAGGGAAGTCGGTGAGGGCGTGAAACCCTGAATCCGGCGCTGCCCCCGCAACTGTGAG
CGGCGAGCCGCTGTCCGTTTCGTGTCACTGACGCGCCGAAGCTGGTTCGGGGATGCGTCGGGAAGGCCAGGGCA
GGGGTGACGACCCGTGAGCCAGGAGACCTGCCTCGACAGATAACGTCCTCCGGCGGGGTGTCCGGTCTGGCCGC
TTGCTCAGCGCGACCGGACAAAAGCGCCCGTGCGCGCTCGACCGCGCGCGTCCCGATCAGCCTCGCCAAAACACC
GGCAGAGGCTTTTCAAAG ATGTTAATTAATATGCATGGTAC

The plasmid pRVChyC-6 (Thanbichler et al., 2007) was cut with MfeI and PacI, and the gBlock segment was inserted into the plasmid by gibson assembly. Next, the resulting plasmid was transformed into E. coli DH5 alpha cells and selected on LB-chlor plates. The resulting chlorR colonies were then screened by PCR for the insert and verified by sanger sequencing (genewiz). The purified plasmid was then transformed into NA1000 cells by electroporation and plated on PYE-chlor plates. The resulting colonies were screened for mChy fluorescence after induction with vanillate.

**JS440**

The plasmid pRVMCS-2 (Kan$^R$) (Thanbichler et al., 2007) was transformed into NA1000 cells via electroporation and selected for on PYE-Kan plates.

**JS441**

This strain was generated by PCR amplifying the last 500bp of the β' RNA polymerase gene into the pYFPC-1 plasmid (Thanbichler et al., 2007). The insert was PCR amplified from the NA1000 chromosome using Betaprime_forward and Betaprime_reverse primers and the plasmid pYFPC-1 (Thanbichler et al., 2007) was PCR amplified by X and Y primers. The plasmid was then DPN 1 treated, and the insert was placed into the plasmid by gibson assembly. Next, the resulting plasmid was transformed into E. coli DH5 alpha cells and selected on LB-spec plates. The resulting specR colonies were then screened by colony PCR for the insert and verified by sanger sequencing (genewiz). The purified plasmid was then transformed into NA1000 cells by electroporation and plated on PYE-spec/strep plates. The resulting colonies were screened for YFP fluorescence.

PCR primers:
Betaprime_forward 5' TAATATGCATGGTGTCGACGAGATCCAGGAGG
Betaprime_reverse 5' ATCTTAAGGTTTCGGCGTCCGAAAGCGC
pYFPC_forward 5' GCTTTCGGACGCCGAAACCTTAAGATCTCGAGCTCCG
pYFPC_reverse 5' GGATCTCGTCGACACCATGCATATTAATTAAGGCGCC
